# snoCLASH Reveals Extensive snoRNA-mRNA Interaction Networks

**DOI:** 10.64898/2025.12.10.693487

**Authors:** Brittany A. Elliott, Gene Yang, Alex K. Choi, Yinzhou Zhu, Wesley R. Freeman, Christopher L. Holley

## Abstract

Small nucleolar RNAs (snoRNAs) are classically defined as guides for modification of ribosomal RNA (rRNA) and small nuclear RNA (snRNA), yet increasing evidence suggests that box C/D snoRNAs also interact with non-canonical transcripts. Systematic discovery of such interactions has been hindered by overwhelming rRNA abundance and technical limitations in RNA-RNA capture. Here, we present snoCLASH, an optimized snoRNA:RNA binding protein (RBP)-based crosslinking, ligation, and sequencing framework that integrates phenol-toluol extraction, polyA enrichment, nuclear fractionation, rRNA depletion, and dual-reference chimeric read analysis to enable transcriptome-scale identification of snoRNA-non-rRNA interactions. Applying this approach reveals thousands of snoRNA-associated mRNA regions spanning coding and regulatory elements and enriched for RBPs linked to epitranscriptomic regulation. Using this framework, we identify high-confidence snoRNA-mRNA interactions and functionally validate *OGA* mRNA as a snoCLASH-discovered target undergoes snoRNA-dependent 2’-*O*-methylation with downstream effects on protein expression. Together, this work establishes snoCLASH as a platform for discovering and validating non-canonical snoRNA targets beyond rRNA.

**GRAPHICAL ABSTRACT:** 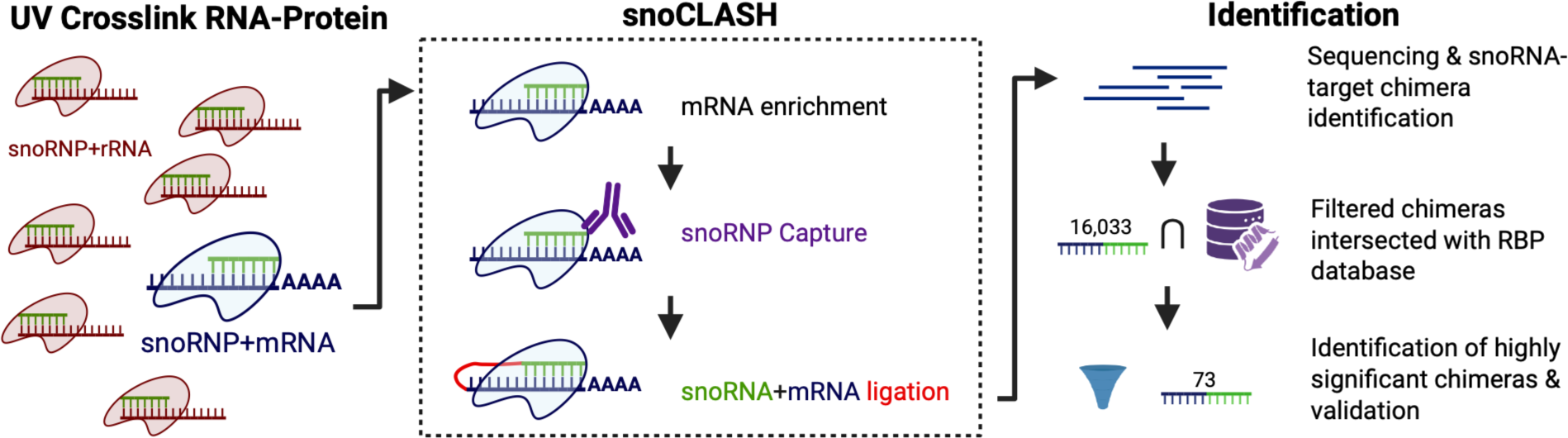

## INTRODUCTION

Small nucleolar RNAs (snoRNAs) are abundant, highly-structured noncoding RNAs best known for guiding 2’-*O*-methylation (Nm) and pseudouridylation on ribosomal RNAs (rRNAs) and small nuclear RNAs (snRNAs) (1, 2). These canonical functions, executed by box C/D and box H/ACA snoRNP complexes, are essential for ribosome biogenesis and translational fidelity (1–4). However, emerging research has begun to reveal a more complex role for snoRNAs, extending beyond their conventional functions. Beyond rRNA modification, snoRNAs have been implicated in pre-mRNA splicing (5–7), RNA stability (8), and stress responses (8–10), and their dysregulation has been linked to cancer (11–16), neurodevelopmental disorders, cardiometabolic disease (9, 17–21), and aging (22, 23). These emerging functions suggest that snoRNAs may act as multifunctional RNA regulators.

A compelling yet incompletely defined hypothesis is that box C/D snoRNAs direct *2-O-* methylation to mRNAs and other non-canonical transcripts. Nm on mRNAs impact translation efficiency, as we and others have demonstrated (24–27). However, unlike rRNA Nm sites, where box C/D snoRNA recognition elements and structural constraints are well established, rules governing snoRNA-mRNA interactions remain undefined. Modeling based on large scale RNA-RNA interaction datasets suggests that snoRNAs interact with targets based on antisense element (ASE) complementarity as well as noncanonical interactions within diverse regions of the snoRNA (28). In parallel, recent experimental approaches using crosslinking and ligation-based sequencing have provided direct evidence of snoRNA interactions with non-ribosomal RNA targets, supporting a broader role for snoRNAs in transcriptome-wide RNA regulation (29-31). Experimental validation of Nm is further challenged by the chemical nature of Nm itself: the modification is small, located on the ribose rather than the nucleobase, chemically inert, and may not be present at high stoichiometry for a given site, making it significantly more difficult to detect than other epitranscriptomic modifications (32).

To address these challenges, we combined optimized snoRNP-based Crosslinking, Ligation, and Sequencing of Hybrids (CLASH) (33) approaches with rRNA-depletion and PolyA-enrichment to physically isolate snoRNA + non-rRNA target chimeras (snoCLASH). We then applied rigorous filtering pipelines and an RNA binding protein (RBP)-centered analysis framework to define the landscape of snoRNA-RNA contacts might influence RBP-RNA interactions, and to identify features that distinguish genuine mRNA interactions from background. Here, we present an integrated analysis of box C/D snoRNA interactomes, including the transcript regions they bind, the RBPs that co-localize with these sites, and the enrichment of modification-associated RBPs that may coordinate or modulate snoRNA-guided Nm on mRNA.

Our findings reveal widespread snoRNA associations with non-ribosomal transcripts, demonstrate that snoRNA target sites overlap with those of known RBPs, and provide a mechanistic framework for how snoRNAs may access and potentially modify mRNAs in vivo. We functionally validate this framework through the identification and validation of O-GlcNAcase (OGA) as a snoRNA-modified transcript, demonstrating that snoRNA-guided Nm can directly influence mRNA translation and downstream protein-dependent signaling pathways. This work expands the regulatory repertoire of snoRNAs and establishes a platform for discovering and validating novel snoRNA-guided Nm targets across the transcriptome.

## MATERIALS AND METHODS

### Buffers, Linkers, Primers, and Enzyme Mixtures

Detailed descriptions are in Supplementary Data.

### Cell Culture and UV Crosslinking

Human embryonic kidney 293T (HEK 293T) cells were cultured to approximately 80% confluency. Cells were then exposed to UV light at 0.45 J/cm^2^ for crosslinking (Boekel Scientific). Following crosslinking, cells were immediately collected for further processing.

### Nuclei Isolation (Fractionation iClip, FriCLIP)

The FriCLIP nuclei extraction protocol was employed for nuclei isolation (34). Cells were washed with ice-cold PBS, detached, and centrifuged at 180 x g for 5 minutes at 4°C. The cell pellet was resuspended in Hypotonic Buffer and incubated on ice for 15 minutes. Subsequent centrifugation at 425 x g for 10 minutes at 4°C separated the cytoplasmic fraction. The pellet was then treated with Lysis Buffer 0.3% and 0.5% sequentially, each followed by incubation on ice for 10 minutes and centrifugation at 950 x g for 10 minutes at 4°C.

### Phenol-Toluol Extraction (PTex)

Crosslinked RNA-Protein complexes were extracted using the PTex method (35). The process involved a three-step extraction using phenol, toluol, and 1, 3-bromochloropropane, followed by ethanol precipitation. The extraction was performed at room temperature with shaking (1, 500 rpm on an Eppendorf ThermoMixer R) and subsequent centrifugation at 20, 000xg for 3 minutes at 4°C. The final pellet was resuspended in 30-50 |_il of distilled water.

### Ribosomal RNA (rRNA) Depletion

rRNA was depleted from the PTex RNA-Protein complexes using a RiboZero kit (Illumina), scaled up according to the quantity of RNA. The rRNA-depleted samples were quantified using a Qubit fluorometer (Thermo) and checked for rRNA depletion with an Agilent bioanalyzer (Agilent).

### Small Nucleolar RNA (snoRNA) Interaction Capture

A protocol modified from eCLIP (36) and qCLASH (37–39) workflows was employed for snoRNA-mRNA interaction capture:

*NOP56 Antibody Binding:* 100 |_iL of Protein A magnetic beads (Thermo) were prepared per sample and bound with 10 |jg of NOP56 antibody (Invitrogen, PA5-78329). The beads were incubated at room temperature for 45 minutes.

*RNA-Protein Fragmentation*: The RNA-Protein complexes were fragmented using RNase I (Thermo) (diluted 1:50), followed by incubation at 37°C for 5 minutes with shaking (1, 200 RPM).

*Immunoprecipitation (IP)*: Fragmented samples were incubated with antibody-bound beads and 11 |_iL of Superasin (Thermo) at 4°C overnight.

*Washing and Phosphorylation*: Post-IP, the beads underwent several washes with different buffers (eCLIP lysis buffer, 1X and 5X PXL buffer, High-Stringency buffer, High Salt buffer, and 1X PNK buffer). The RNA was then phosphorylated using a T4 PNK Mixture (10X PNK Buffer, Superasin, 100 mM ATP, RNase free Water, T4 PNK) at 10°C for 40 minutes.

*Intermolecular Ligation*: The RNA was ligated to a 3’ RNA linker using a T4 Ligation Mixture (10X T4 RNA ligase buffer, 50% PEG-8000, 2 M KCl, Superasin, 100 mM ATP, RNase Free Water, T4 RNA ligase 1 (NEB)) at 4°C overnight.

*Dephosphorylation and 3’ Ligation*: Subsequent dephosphorylation and 3’ ligation were performed using an Antarctic Phosphatase Mix (NEB) (10X Antarctic Phosphatase Buffer, Superasin, RNase Free Water, Antarctic Phosphatase) at 10°C for 40 minutes with periodic shaking. 10 uM of a modified miRCat-33 linker was added with 80 uL 3’ Linker Ligation Mix (T4 RNA ligase buffer, 50% PEG8000 (16 uL), Superasin (4 uL), and T4 RNA ligase 2, truncated KQ (NEB, 4 uL) overnight at 16C with shaking every 2 minutes for 15 seconds at 1, 000 RPM.

*Wash and Elute from Beads:* Beads were washed three times with 1X PNK Buffer. 100 |_iL Elution Buffer was added to the beads and incubated at room temperature for 15 minutes at 1, 400 RPM. The supernatant was transferred to a new tube and the steps were repeated for complete elution.

*Proteinase K Treatment*: A Proteinase K (Thermo) mixture (5X Proteinase K Buffer, Proteinase K, RNase Free Water) was prepared and incubated at 37°C for 20 minutes. This mixture was then added to the supernatant from the previous step and incubated for an additional 20 minutes at 37°C.

*TRIzol Extraction*: RNA was extracted using TRIzol (Thermo) according to the manufacturer’s instructions and eluted in 10 |_iL water.

*5’ Phosphorylation and Linker Ligation*: RNA was phosphorylated using a T4 PNK Mixture, followed by ligation to a 5’ RNA linker using a T4 RNA ligase 1 (NEB) -based ligation mixture.

This was incubated overnight at 16°C.

*Reverse Transcription and PCR Amplification*: Reverse transcription was conducted using a mix of 5X Superscript RT buffer, DTT, RNase Out, and Superscript III (Thermo), followed by PCR amplification with Q5 high-fidelity mastermix (NEB) and Illumina sequencing primers.

*Sample Cleanup and Precipitation*: PCR products were cleaned of salts and contaminates using NUC-away columns (Thermo).

*Gel Purification and DNA Purification*: PCR products were run on a 4% agarose e-gel (Thermo) with SybrGold (Thermo), and fragments above 200 bp were excised and purified.

*Bioanalyzer Analysis*: The final purified samples were analyzed using an Agilent Bioanalyzer (Agilent) with a High Sensitivity DNA chip to assess the quality and size distribution of the captured RNA-DNA complexes.

### RNA-sequencing

Sample libraries generated to capture snoRNA-mRNA targets as described above were multiplexed and sequenced with an Illumina MiniSeq (150 nt Single-Read) and NovaSeq 6000 (75 nt Paired-End).

### Chimeric Read Analysis (ChiRA)

Reads were mapped and analyzed for snoRNA-mRNA chimeras using the ChiRA pipeline (40, 41) to genome build hg38/GRCh38. A custom hg38 reference was generated to be compatible with the pipeline and to include predicted as well as canonically known snoRNA species. ChiRA’s dual-reference alignment strategy mapped each read simultaneously to the snoRNA reference set and the transcriptome/genome, retaining only high-confidence hybrid assignments. Raw chimeras were first collapsed by read sequence to remove PCR and sequencing duplicates. For each unique chimera, ChiRA-reported alignment information was retained for downstream filtering, including alignment start and end positions, transcript/gene annotations, and snoRNA identity. Because the ChiRA dual-reference annotation framework assigns interacting fragments to annotated transcript features, chimeras involving snoRNA-snoRNA interactions could not be explicitly resolved as direct snoRNA targets. In these cases, interactions involving intronic snoRNAs were typically assigned to the corresponding host-gene intronic region or host non-coding RNA transcript rather than the embedded snoRNA itself.

### Post Data Processing

To eliminate chimeras arising from low-complexity or false sequencing reads, hybrids containing ≥8 identical consecutive nucleotides making up greater than >50% of the identified transcript on either side of the junction were removed. Chimeras mapping to mitochondrial rRNA or nuclear rRNA were also removed to avoid inflating non-informative high-copy species. Transcript variants and overrepresented high-copy species were also removed from the POSTAR3 database to reflect this trimming of test data. AI tools were used to write code for processing and plotting some of the computational data.

### RNA-binding protein (RBP) comparison

Transcript regions captured with snoRNAs were intersected with the POSTAR3 database of RNA-binding protein-RNA interactions (42) using bedtools intersect (43) function to detect regions intersecting by one nucleotide or more. RBP peaks with a score of three or more were included in the intersection.

To determine whether certain RBPs were statistically enriched at snoRNA-target sites, each RBP’s POSTAR3 peaks were subjected to a transcript-aware bedtools shuffle (43). Peaks from each RBP (n=100) were randomly reassigned (10, 000 permutations) to transcript regions matching their original annotation class (e.g., CDS peaks shuffled only within CDS), and intersects with snoRNA-target intervals were recomputed at each iteration. RBPs whose observed intersect count exceeded the 99th percentile of their permutation-derived null distribution (α = 0.01) were classified as significantly enriched.

To avoid false positives driven by extreme or low-quality peak scores, enriched RBPs were further filtered by requiring an average POSTAR3 peak score ≥20 after exclusion of both lower and upper 25% outliers. This produced a stringent high-confidence RBP set enriched at snoRNA-target sites.

To evaluate whether RNA-modification-associated RBPs (e.g., writers, readers, and erasers of epitranscriptomic modifications) were enriched among the RBPs identified as significantly overlapping snoRNA-target regions, we applied a hypergeometric model. Within this background set (POSTAR3 database; N=221), the number of RBPs annotated as RNA-modification-associated was recorded (K=25). The subset of RBPs found to be significantly enriched in the permutation-based analysis (n=28) was treated as the observed “draw, ” and the number of RNA-modification RBPs within this enriched set (x=12) was used as the test statistic.

The probability of observing x or more RNA-modification-associated RBPs in the enriched set, given random sampling without replacement, was computed using the survival function of the hypergeometric distribution:

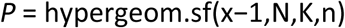

This test evaluates whether the observed number of RNA-modification RBPs in the enriched set exceeds what would be expected under random sampling from the background distribution. A p-value ≤ 0.05 was considered statistically significant.

### IntaRNA

Predicted snoRNA-target interactions were evaluated using IntaRNA (v3) in pairwise mode (44). Query and target sequences derived from snoCLASH chimeras were processed in chunked FASTA files to improve computational efficiency. Interactions were constrained to a maximum interaction length of 30 nucleotides (--intLenMax=30), corresponding to the typical region between the conserved box C and box D elements of box C/D snoRNAs, which contains the antisense guide sequence responsible for target recognition. IntaRNA was run using the parameters --seedBP=5 and --seedNoGU to require a minimum 5 bp seed interaction without G:U wobble pairing. Accessibility calculations were restricted to 30 nt windows for both query and target sequences (--qAccL=30 --qAccW=30 --tAccL=30 --tAccW=30). Only the top-scoring interaction per pair was retained (--outNumber=1).

### RNA Extraction for qPCR Assays

RNA was extracted from 293T cells and mouse heart usng TRIzol (Thermo), per manufactuer’s directions. cDNA was synthesized with 1 ug total RNA with oligo dT as a reverse primer and Superscript III as the reverse transcriptase according to manufacturer instruction (Thermo). All qPCR analyses were performed and analyzed with technical duplicates, using PowerSybr (Thermo) and an ABI StepOnePlus Real-Time PCR System.

### Reverse Transcription at Low dNTP concentrations followed by PCR (RTL-P)

RTL-P was performed with described modifications (27, 45) to the originally published method (46). cDNA was generated from 1 ug of total RNA, with oligo-dT RT priming, under two conditions: (1) standard Superscript III (Thermo) reverse transcriptase reaction (1 mM dNTP mix), and (2) under the same conditions except low dNTP concentration (0.1 mM dNTP mix). qPCR was then performed in 20 |_iL reactions with Power SYBR (Thermo), 0.25 of Nm sensitive (both 5’ to the Nm site) and Nm insensitive (both 3’ to the Nm site) forward and reverse primers, and 2 |_iL of each cDNA with a hot start (95 C; 10’) and 40 cycles of 95 C (15 s) and 60 C (1’). Relative quantification of *OGA* from each cDNA reaction (both high and low dNTP) was first normalized to an internal reference gene (*RPLP0*/*Rplp0*) to correct for any differences in total RNA input (standard Pfaffl method) (47). The ratio of low/high dNTP product was then calculated for each condition, to normalize the low dNTP result to the amount of *OGA/Oga* mRNA in each sample. Finally, this value could be compared between experimental (KO) conditions as a relative quantity (RQ). This method accounts for changes in transcript methylation relative to transcript abundance in normal conditions.

### Animals

*Rpl13a* snoRNA KO mice have previously been described (17, 27). Their humane and ethical use was approved by the Duke Institutional Animal Care and Use Committee (IACUC), Protocol Registry Number A219-21-11-24.

### Western Blotting

Mouse heart lysates were prepared in RIPA buffer (Sigma R0278) with cOmplete™ Protease Inhibitor Cocktail (Roche 4693116001). 293T clones were collected in hypotonic lysis buffer supplemented with protease inhibitor and collected following a previously published method (48). Protein concentrations were determined using a BCA assay (ThermoFisher 23225).

Proteins were separated on 4-15% SDS-PAGE gels and transferred to nitrocellulose membranes. Immunoblotting was performed with the following antibodies: anti-O-GlcNAcase (OGA) (Novus Biologicals, NBP2-32233, 1:1000). Images were captured using a ChemDoc XRS + system (Bio-Rad). Protein expression levels were normalized to total protein imaging and analyzed using ImageLab software (Bio-Rad).

### Statistical Analyses

Statistical analyses were performed using GraphPad Prism (v9 or later) unless otherwise specified. For comparisons between two groups, statistical significance was assessed using two-tailed unpaired Student’s *t*-tests. Data are presented as mean ± standard error of the mean (SEM), with individual data points shown where applicable. Group comparisons for these assays were performed as indicated in the corresponding figure legends. Spearman rank correlation coefficients (ρ) were calculated using Python (SciPy v1.x) to assess monotonic relationships between variables. Correlation analyses were performed to evaluate the association between RQ or RTL-P measurements and protein expression levels. Two-tailed P-values were calculated for all correlation analyses. Exact P-values and correlation coefficients are reported in the figures.

## RESULTS

### Optimized snoCLASH workflows enrich for non-ribosomal snoRNA-target chimeras

CLASH has previously been shown by multiple groups to predominantly recover snoRNA-rRNA hybrids due to the vast stoichiometric excess of rRNA in cells and the strong affinity between snoRNPs and their canonical rRNA substrates. This is illustrated in the schematic of a typical CLASH workflow (**Fig. 1A Column I**). To overcome this limitation, we have developed two modified snoCLASH approaches specifically designed to reduce rRNA abundance and increase the recovery of non-ribosomal snoRNA-RNA interactions. NOP56 was selected as the bait protein for snoRNP immunoprecipitation because it maintains stable physical contact with box C/D snoRNAs throughout RNP assembly and function, in contrast to the 2’-*O*-methylase Fibrillarin (FBL), which interacts transiently with snoRNA-target complexes only during catalysis. In addition, a high-performing NOP56 antibody was available for efficient and reliable immunoprecipitation.

**Figure 1.**
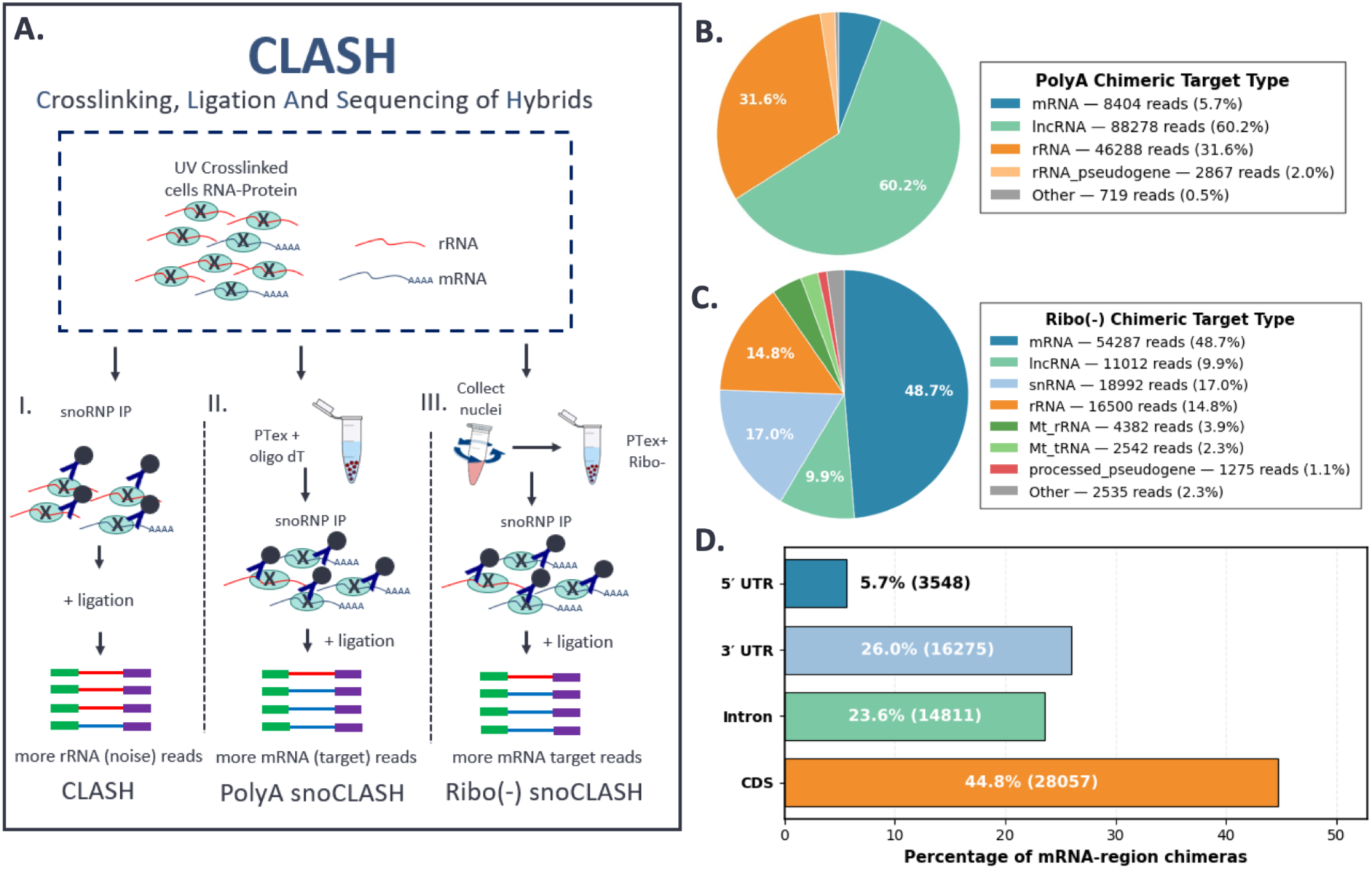
ChiRA Analysis of Crosslinked RNA-Protein Complexes Reveals Non-rRNA Targets. **A.** Overview of the snoRNA-target CLASH workflow. UV-crosslinked RNA-protein complexes are subjected to snoRNP immunoprecipitation (IP) followed by RNA ligation to capture snoRNA-target RNA hybrids. Three enrichment strategies are shown: (I) Standard CLASH, which yields predominantly rRNA-derived hybrids (“noise” reads); (II) PolyA snoCLASH, in which phenol-toluol extraction (PTex) coupled with oligo-dT selection increases recovery of non-rRNA targets; and (III) Ribo-snoCLASH, in which nuclei are collected and rRNA is depleted using PTex + Ribo-treatment prior to snoRNP IP and ligation, enriching for non-ribosomal RNA hybrids. **B.** Distribution of RNA biotypes recovered in PolyA snoCLASH. The majority of captured hybrids map to IncRNA (60.2%), with additional reads mapping to rRNA (31.6%), transcripts, pseudogenes, and other minor RNA classes. **C.** RNA biotype distribution in Ribo-snoCLASH. rRNA depletion markedly increases recovery of mRNA-derived hybrids (48.7%). **D.** Distribution of snoRNA-target chimeras across mRNA regions in PolyA and Ribo(-) snoCLASH libraries. Chimeras most frequently map to coding sequences (CDS), followed by 3’ UTRs and introns, with relatively few detected in 5’ UTRs.

A key innovation in both optimized workflows was the use of phenol-toluol extraction (PTex) to selectively purify RNA molecules that were physically crosslinked to proteins (35). By enriching for protein-associated RNA while removing the majority of free or background RNA species, PTex greatly reduced nonspecific carryover and enabled downstream application of specialized reagents, such as oligo-dT for polyA selection or Ribo(-) depletion reagents, that cannot be effectively applied to bulk RNA in amounts required for effective immunoprecipitation.

Chimeric reads generated from these optimized workflows were then processed with the ChiRA chimeric-read analysis framework, which maps, annotates, and filters RNA-RNA hybrids with high specificity, enabling transcriptome-wide identification of snoRNA-target chimeras (41). Despite strategies to reduce rRNA capture, snoCLASH recovered 33 unique chimeras with box C/D snoRNAs and 8 H/ACA snoRNAs engaged with their canonical rRNA targets, with chimeric reads overlapping known or predicted modification sites (**Supplementary Table 1**) (49, 50).

In PolyA-snoCLASH (**Fig. 1A Column II, Supplementary Fig. 1**), PTex-purified RNA-protein complexes were subjected to oligo-dT selection to enrich mRNAs before NOP56 immunoprecipitation. Efficient enrichment of polyadenylated RNA and NOP56-associated complexes following crosslinking and PTex extraction was confirmed by immunoblot analysis (**Supplementary Fig. 2**). This strategy decreased rRNA contamination and substantially increased the recovery of snoRNA hybrids with non-ribosomal RNAs, with the majority of chimeric reads mapping to lncRNAs (**Fig. 1B**).

In Ribo(-) snoCLASH (**Fig. 1A Column III, Supplementary Fig. 3**), two additional steps were added to further reduce rRNA. First, nuclei were isolated immediately after UV crosslinking (34), removing the vast majority of cytoplasmic ribosomes and thereby dramatically lowering rRNA background. This nuclear-enriched fraction was ideal for targeting snoRNA interactions because box C/D snoRNAs primarily localize to nucleoli, where they encounter RNA targets. Second, PTex-purified nuclear complexes were treated with Ribo(-) depletion reagents, which efficiently removed remaining rRNA and pre-rRNA species. This enabled enhanced detection of non-rRNA targets within the nuclear compartment, with almost half of chimeric reads mapping to mRNA (and/or pre-mRNA) (**Fig. 1C**). Ribo(-) snoCLASH also captured snoRNA-snRNA interactions, as expected.

When PolyA-snoCLASH and Ribo(-) snoCLASH datasets were analyzed together, snoRNA-associated mRNA and pre-mRNA targets were distributed broadly across transcript regions, including coding sequences (CDS; 44.8%), 3’ untranslated regions (3’ UTR; 26%), introns (23.6%), and 5’ UTRs (5.7%) (**Fig. 1D**). This wide distribution demonstrates that snoRNAs engage non-ribosomal RNAs throughout both coding and regulatory regions, revealing a landscape of interactions that is distinct from findings by conventional methods.

### Many high-confidence snoRNA-target regions intersect with RBP-bound sites

To distinguish bona fide snoRNA-RNA interactions from crosslinking background, we applied a stringent post-processing pipeline involving deduplication, removal of repetitive chimeras (≥8 identical consecutive nucleotides), removal of high-copy species (SNORD3, nuclear and mitochondrial rRNA, and mitochondrial RNA), and filtering of assignments to single representative target intervals (**Fig. 2A**; detailed workflow diagrams in **Supplementary Fig. 4**). After filtering, 16, 033 unique snoRNA-target regions remained (**Fig. 2A, B**). We hypothesized that if snoRNAs guide functionally relevant modifications to RNA, target sites might preferentially localize to transcript regions already engaged by RBPs, particularly those known to read, write, or interpret RNA modifications (51–54), as RNA modifications are non-randomly positioned (55–57) at sites that regulate transcript stability, translation, and fate (58). Intersecting these target sites with the POSTAR3 database of experimentally mapped RBP-RNA interactions (42) revealed that 6, 450 of the 16, 033 unique snoRNA targets (40.2%) overlapped at least one RBP peak (POSTAR3 RBP Score > 20) by >1 nt (**Fig. 2B**), with the majority mapping to mRNA (**Fig. 2C**).

**Figure 2.**
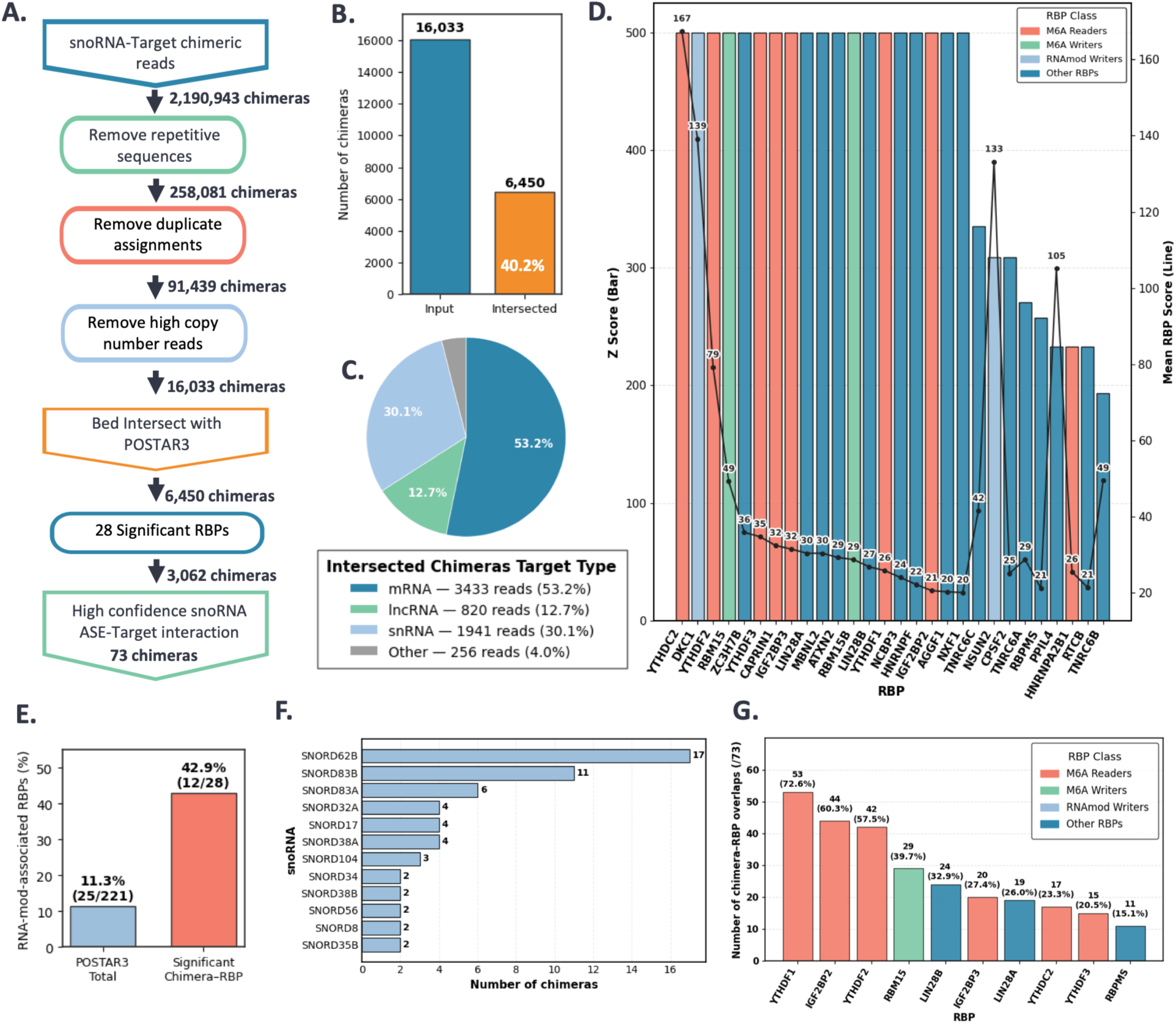
Identification and enrichment of RNA-binding proteins at snoRNA-target chimera sites. **A.** Workflow for filtering snoRNA-target chimeras prior to RBP intersection. Duplicates, low-complexity reads (>8 identical consecutive nucleotides), and high-copy RNAs (SNORD3, nuclear and mitochondrial rRNA, and mitochondrial RNA) were removed. High-confidence regions were intersected with POSTAR3 RBP binding peaks. B. Overlap between filtered chimeras and POSTAR3 peaks. Of 16, 033 unique chimeras, 6, 450 (40.2%) intersected at least one RBP peak (score > 20) by >1 nucleotide. **C.** Genomic distribution of RBP-overlapping targets, enriched in mRNA (53.2%), snRNA (30.1%), and IncRNA (12.7%). **D.** Identification of significantly enriched RBPs by permutation analysis. RBP peaks were shuffled 10, 000x within matched transcript regions to generate null distributions; RBPs exceeding the 99th percentile (a = 0.01) were retained and further filtered by IQR-trimmed mean score > 20. **E.** RNA-modification-associated RBPs are enriched among significant hits (42.9% vs. 11.3% background; 3.8-fold enrichment, *p =* 2.85 × 10^6^; hypergeometric test). **F.** snoRNAs recurrently identified within the 73 high-confidence snoRNA chimeric interactions, showing snoRNA species detected in two or more unique chimeric interactions. Bar labels indicate chimera count. **G.** Top 10 RNA-binding proteins (RBPs) overlapping high-confidence snoRNA ASE-target chimera regions, ranked by the number of overlapping chimeras. Bar labels indicate the number and percentage of the total 73 high-confidence interactions associated with each RBP.

### Permutation analysis identifies RBPs significantly enriched at snoRNA-target sites

To identify RBPs that associate with snoRNA-bound regions more frequently than expected by chance, we first performed a permutation-based enrichment analysis across all POSTAR3 RBPs. For each RBP, all its binding peaks were shuffled 10, 000x into transcript regions matching their original annotation class (e.g., intron to intron), and snoRNA-target intersects were recomputed at each iteration to generate an empirical null distribution (**Supplementary Fig. 4**). RBPs whose observed intersect counts exceeded the 99th percentile of their shuffled distributions (α = 0.01) were classified as significantly enriched.

After this initial statistical filtering, we further refined the significant RBP set by requiring an average POSTAR3 binding site score ≥ 20. To minimize bias from extreme or low-quality peaks, average binding site scores were computed after restricting values to the interquartile range (IQR; 25th-75th percentiles) (59), and RBPs with an IQR-filtered mean score ≥ 20 were retained. This two-tiered approach (statistical enrichment followed by binding-score refinement) yielded a stringent, high-confidence set of 28 RBPs most likely to interact functionally with snoRNA-bound target regions (**Fig. 2D**). Corresponding Z-scores are shown by the bar graphs and mean POSTAR3 binding site scores (“mean RBP scores”) are shown by the line plot.

### RNA-modification-associated RBPs are enriched at snoRNA-target sites

After defining the high-confidence set of significantly enriched RBPs, we next examined whether specific functional classes of RBPs were overrepresented. Of the 221 RBPs present in the filtered POSTAR3 dataset, 25 are classified as epitranscriptomic RNA modification proteins (11.3% of 221). However, among the 28 significantly enriched RBPs derived from our permutation analysis, 42.9% (12/28) belonged to this category (**Fig. 2E**), a striking overrepresentation relative to background. To quantify this enrichment, we applied a hypergeometric model comparing the observed number of RNA-modification RBPs within the significant set to the number expected by random sampling from the POSTAR3 RBP population. This analysis revealed a 3.8-fold enrichment of RNA-modification RBPs (p = 2.85 × 10^−6^), demonstrating that RNA-modifying enzymes and readers are strongly and non-randomly associated with snoRNA-target interaction sites.

### Selection of high-confidence snoRNA-target interactions with significant RBPs

To further refine the biological relevance of these overlaps, we focused on snoRNA-target chimeras supported by predicted complementarity within known or predicted snoRNA antisense regions (60) (IntaRNA ΔG < -10 kcal/mol) (44) and intersecting significantly enriched RBPs identified through the permutation-based analysis. This filtering strategy yielded 73 high-confidence snoRNA antisense element (ASE)-target interactions associated with enriched RBP binding regions (**Supplementary Table 2**). We next examined the snoRNA composition of this high-confidence set and observed representation of 26 different box C/D snoRNAs, with 12 snoRNAs appearing in two or more independent chimeric interactions (**Fig. 2F**). Among the 73 high-confidence snoRNA ASE-target interactions, chimeras were not uniformly distributed across the 28 significantly enriched RBPs identified by permutation analysis, but instead clustered within a subset of RBPs (**Fig. 2G**). Notably, seven of the top 10 RBPs most frequently overlapping these high-confidence chimeric regions are known m6A readers (YTHDF1, IGF2BP2, YTHDF2, IGF2BP3, YTHDC2, and YTHDF2) or an m6A writer (RBM15).

### SNORD32A guides a candidate Nm site on the OGA transcript

Among the 73 high-confidence snoRNA ASE-target interactions intersecting significantly enriched RBPs, we selected the *SNORD32A*-*OGA* chimera for targeted validation. This interaction was prioritized because *SNORD32A* is of particular biological interest given its established role in mitochondrial ROS production and metabolic stress responses (9, 17, 20, 27), and our laboratory has existing genetic and experimental tools enabling direct functional interrogation of this snoRNA. The *SNORD32A-OGA* chimera site overlaps with a site of known interaction between OGA and RBP IGF2BP2, which is a known m6A reader and the second most frequently overlapping RBP within the high-confidence dataset (**Fig. 2G**). Furthermore, *OGA* encodes the major enzyme responsible for removal of protein O-GlcNAc modification, linking this candidate interaction to pathways previously associated with ROS and *SNORD32A* biology (61). IntaRNA modeling of the fragment captured in snoCLASH predicts a stable interaction between the *SNORD32A* 5’ antisense element (ASE) and an *OGA* exon region, with a calculated free energy of -12.54 kcal/mol (**Fig. 3A**). The modeled duplex predicts *SNORD32A*-guided Nm modification of an adenosine on OGA mRNA, in a manner consistent with canonical box C/D snoRNA-guided Nm deposition (5 nucleotides upstream of the snoRNA D-box).

**Figure 3.**
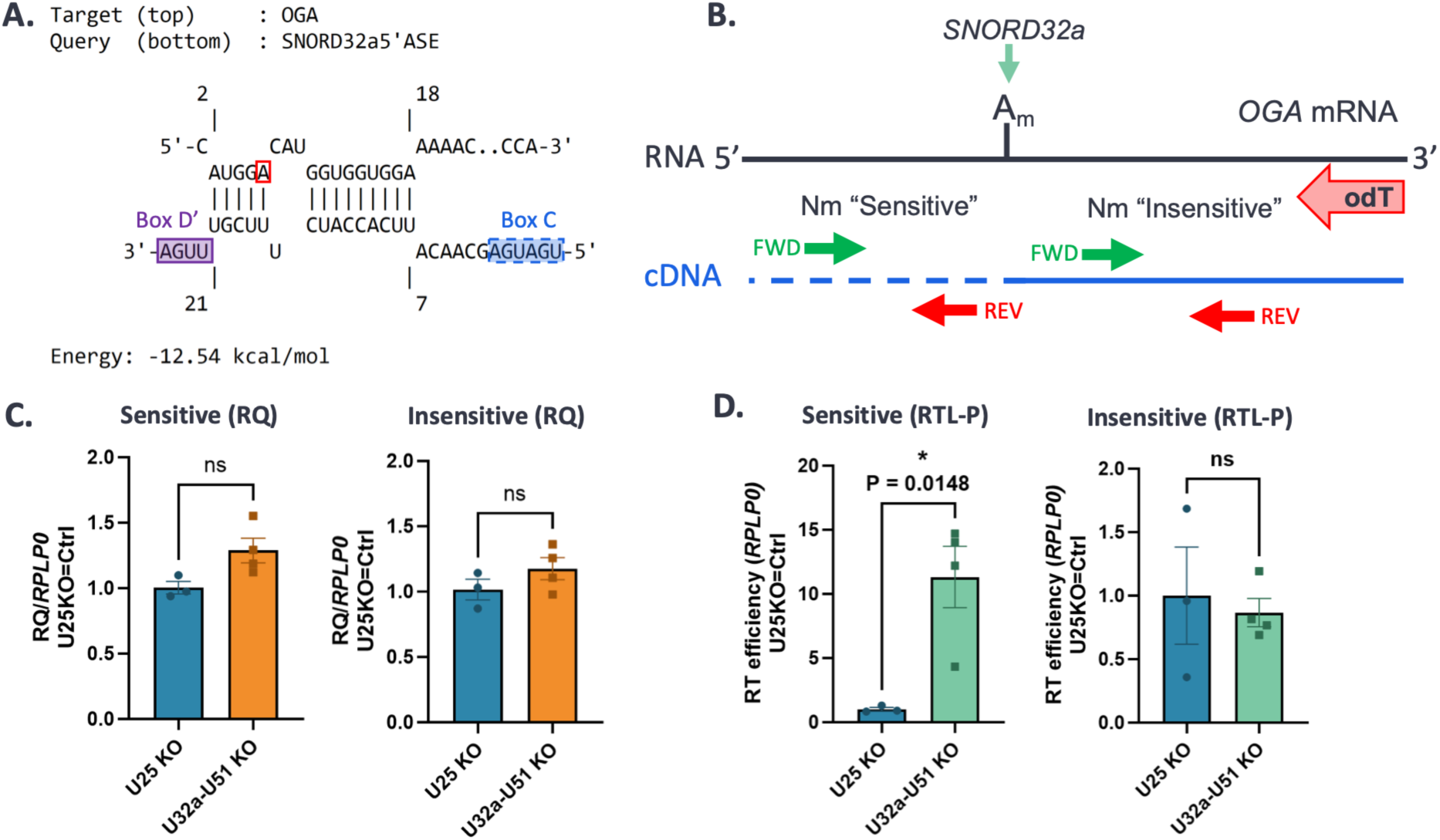
*SNORD32A* interacts with the OGA transcript and guides a candidate Nm site validated by methylation-sensitive reverse transcription. **A.** IntaRNA interaction model between *SNORD32A’s* 5’ antisense element (ASE) and the OGA transcript fragment captured by snoCLASH. The predicted duplex (AG = -12.54 kcal/mol) positions a central adenosine consistent with box C/D snoRNA-guided Nm targeting. **B.** Schematic of the *OGA* transcript and experimental design. Nm-sensitive primers anneal 5’ of the predicted Nm site and report on 2’-O-methylation using Reverse Transcription at Low dNTPs (RTL-P). Nm-insensitive primers anneal 3’ of the site and are unaffected by Nm. Oligo-dT was used to prime reverse transcription under high- and low-dNTP conditions. **C.** Relative quantification (RQ) of OGA transcript abundance in WT 293T cells and CRISPR knockout clones lacking control snoRNA *SNORD25* or test snoRNAs *SNORD32A* and *SN0RD51.* No significant differences were observed in *OGA* abundance for either primer set, confirming that loss of *SN0RD32A/SN0RD51* does not alter transcript levels. **D.** RTL-P analysis of the predicted Nm site. Using Nm-sensitive primers, *SN0RD32A/SN0RD51* knockout clones show significantly increased RT efficiency (reduced stalling), indicating loss of Nm at the predicted site (P = 0.0148, ns = not significant, t-test). Non-significant differences were observed using Nm-insensitive primers.

To evaluate whether *SNORD32A* directs Nm on *OGA*, we performed Reverse Transcription at Low deoxy-ribonucleoside triphosphate (dNTP) concentrations followed by polymerase chain reaction (RTL-P) (45, 46), using primers sensitive and insensitive to the predicted Nm site (**Fig. 3B**). Because *SNORD51* has a similar ASE that also guides Nm modification at *SNORD32A*’s canonical 28S:Am1511 site, we used 293T clones with both of these snoRNAs knocked out to ensure complete loss of *SNORD*-guided activity from this ASE (27). We then compared these CRISPR knockout clones lacking *SNORD32A* and *SNORD51* to 293T clones lacking the unrelated snoRNA *SNORD25* as a control. Relative quantification (RQ; at normal dNTP levels) showed no change in *OGA* transcript abundance between *SNORD25* control KOs and *SNORD32A*/*SNORD51* KOs for either primer set (**Fig. 3C**), indicating that loss of the snoRNAs does not alter *OGA* transcript levels.

In contrast, the RTL-P assay demonstrated a significant increase in RT efficiency in *SNORD32A/SNORD51* KO cells using the Nm-sensitive primer set at low dNTP concentrations, consistent with the loss of Nm modification (**Fig. 3D**). No difference was observed with the Nm-insensitive primers. These results support the presence of an Nm modification within the *SNORD32A*-predicted region of OGA in WT cells and show that this modification is lost in *SNORD32A*/*SNORD51* KO cells.

### Loss of *SNORD32A* increases OGA protein abundance in human cells and mouse heart

Given our validation that *SNORD32A* guides an Nm site within the *OGA* coding sequence, and reports from our group and others showing that Nm within coding regions can influence translation by modulating ribosome progression, we next examined whether loss of *SNORD32A* affects OGA protein expression. Western blot analysis of *SNORD32A*/*SNORD51* KO clones revealed a >2-fold increase in OGA protein abundance compared to *SNORD25* KO controls (**Fig. 4A-B**). This increase occurred despite unchanged OGA transcript levels in the same cell lines (**Fig. 3C**), consistent with a model in which the loss of Nm in an mRNA coding region enhances translation. To further examine the relationship between transcript abundance, methylation-dependent RTL-P signal, and protein expression, we performed correlation analyses, which revealed a stronger association between RTL-P measurements and OGA protein levels than with transcript abundance (**Supplementary Fig. 5**).

**Figure 4.**
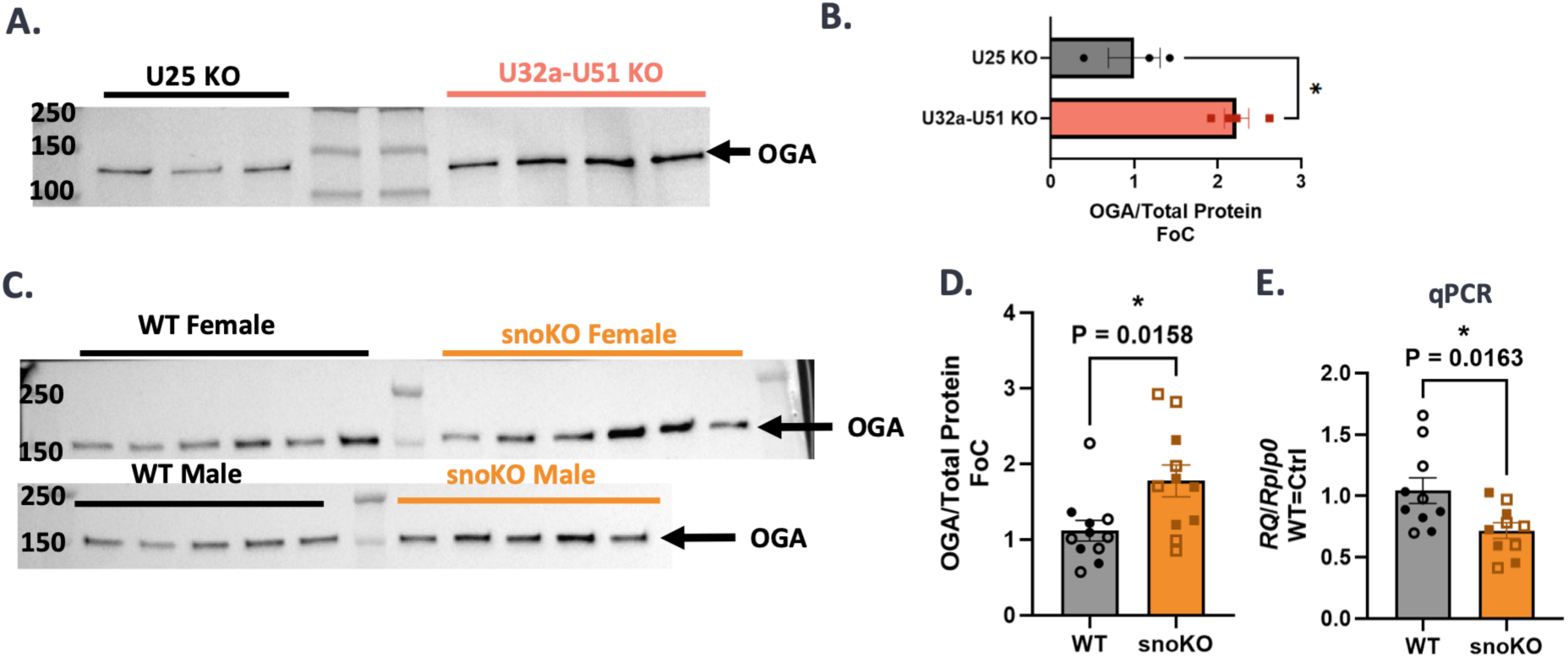
Loss of *SNORD32A* alters OGA protein abundance. **A.** Western blot analysis of OGA protein in human 293T cells lacking control snoRNA U25 or the test snoRNAs SNORD32A/SNORD51. **B.** Quantification of blot **(A.)** showing a significant increase in OGA protein levels in SNORD32A/SNORD51 KO cells compared to U25 KO controls (P = 0.0360, t-test; FoC = fold-change). **C.** Western blot analysis of OGA protein in mouse hearts from WT and *Rpll3a* snoRNA knockout (snoKO) animals (males [closed points] and females [open points]). The snoKO mice lack all four flp/13o-encoded box C/D snoRNAs, including *Snord32A.* **D.** Quantification of blot **(C.)** demonstrating significantly increased OGA protein in snoKO hearts relative to WT controls (P = 0.0158, t-test). **E.** qPCR analysis of OGA transcript levels in WT and snoKO heart tissue. A modest but significant decrease in OGA mRNA was observed in snoKO hearts (P = 0.0163, t-test).

To determine whether this regulatory effect extends in vivo, we evaluated OGA protein levels in heart tissue from *Rpl13a* snoRNA knockout (snoKO) mice, which lack all four *Rpl13a*-encoded box C/D snoRNAs, including *Snord32A*. OGA protein abundance was significantly elevated in snoKO hearts compared to WT controls (**Fig. 4C-D**). Interestingly, *OGA* mRNA levels were modestly but significantly decreased in snoKO animals (**Fig. 4E**), reinforcing that the observed protein increase is not driven by transcription and is more consistent with a post-transcriptional mechanism (27).

## DISCUSSION

In this study, we developed and applied snoCLASH approaches to overcome the persistent technical obstacles that have limited discovery of snoRNA-mRNA interactions in vivo. Traditional CLASH protocols are dominated by snoRNA-rRNA hybrids, reflecting both the overwhelming excess of rRNA and the strong affinity of box C/D snoRNPs for their canonical substrates. Any method seeking to reveal non-rRNA snoRNA targets must overcome three major barriers: (i) physical masking by abundant rRNA, (ii) non-specific RNA background that obscures bona fide interactions, and (iii) the intrinsic difficulty of validating Nm because of its subtle and chemically inert ribose-linked structure. By strategically integrating PTex, polyA enrichment, and nuclear isolation followed by rRNA depletion, we substantially reduced rRNA background and enabled physical capture of snoRNA-mRNA interactions across the transcriptome. Another advance of this work is the application of the ChiRA pipeline, previously used primarily for miRNA-target chimera analysis, to snoRNA-target discovery. ChiRA’s dual-reference alignment strategy is designed to handle CLASH data, enhancing discovery of non-coding RNA-RNA hybrids (41). These optimized workflows revealed thousands of snoRNA-target chimeras distributed across coding and regulatory regions, demonstrating that box C/D snoRNAs engage a much broader spectrum of transcripts than canonical models predict. A caveat of this dual-reference strategy is that snoRNAs were isolated into the dedicated snoRNA reference and excluded from the secondary transcriptome reference used for target assignment, preventing explicit annotation of snoRNA-snoRNA interactions and instead frequently assigning interactions involving intronic snoRNAs to the corresponding host-gene intron or host non-coding RNA transcript.

A major conceptual advance emerging from our study is the demonstration that snoRNA-interacting mRNA sites are non-randomly associated with RNA-binding proteins linked to post-translational RNA modification pathways. Using POSTAR3 intersect analysis and a hypergeometric testing model, we found that RBPs associated with epitranscriptomic writers, readers, and erasers of chemical RNA modifications are disproportionately enriched at snoRNA-bound target regions. This observation suggests that mRNA-bound snoRNAs operate within biochemical environments where RNA modification enzymes are already active, raising the possibility that snoRNAs may collaborate with or recruit these factors to modulate target RNA fate. The 3.8-fold enrichment of RNA-modification-associated RBPs among significant hits provides strong statistical support for this model and places snoRNA-mRNA interactions within an epitranscriptomic regulatory context.

Our integrated analysis further narrows this landscape to a subset of high-confidence snoRNA-mRNA interactions supported by multiple independent criteria, including predicted antisense complementarity (IntaRNA ΔG < -10 kcal/mol), detection across optimized snoCLASH datasets, and overlapping interaction sites with statistically enriched RBPs. The convergence of these features indicates that these chimeras are unlikely to arise from background crosslinking or noise. The presence of these high-confidence interactions suggests that snoRNA engagement with mRNAs reflects a conserved and potentially regulated mechanism by which box C/D snoRNAs can access and influence non-ribosomal transcripts.

Our functional validation of the *SNORD32A-OGA* interaction provides direct evidence of such regulatory capacity. RTL-P assays revealed that *SNORD32A* guides Nm modification within the *OGA* mRNA coding region, and deletion of *SNORD32A/SNORD51* prevents this modification. Importantly, removal of this Nm increases OGA protein abundance in both human cells and mouse heart tissue. Several studies have shown that Nm within coding regions can influence translation by interrupting ribosome elongation, and our data are consistent with this model (24–27). Loss of *SNORD32A*-guided Nm on *OGA* mRNA likely increases translation efficiency in the same way, resulting in elevated OGA protein levels.

While our snoCLASH framework significantly expands the detectable interactome, several limitations merit consideration. First, snoCLASH captures RNA-RNA duplexes present at the point of crosslinking but does not directly measure Nm modification; integration with base-resolution Nm mapping technologies will be essential for correlating snoRNA interactions with Nm modification across the full set of identified targets. Second, transient or low-abundance interactions may remain underrepresented despite PTex, polyA enrichment, and rRNA depletion. Third, although permutation-based RBP enrichment highlights potentially important regulatory regions, biochemical validation will be necessary to determine whether these RBPs act collaboratively, competitively, or independently of snoRNA-guided Nm.

Despite these limitations, our work establishes a framework for discovering non-canonical snoRNA targets and identifying candidate regulatory Nm sites on mRNAs. The snoCLASH approach enables transcriptome-wide identification of snoRNA-RNA interactions and can be readily integrated into existing CLIP-based analysis pipelines. By coupling physical capture of RNA-RNA hybrids with RBP-centric computational frameworks, this method extends the interpretability of CLIP datasets to include direct RNA-RNA interactions. Integration of snoCLASH with RBP occupation analysis, antisense complementarity modeling, and functional validation creates a roadmap for dissecting snoRNA-mRNA networks and their mechanistic consequences. More broadly, our findings highlight snoRNAs as active participants in epitranscriptomic regulation of mRNA, capable of modulating translation and downstream biological pathways through site-specific Nm.

In conclusion, our findings support and experimentally confirm our initial hypothesis: box C/D snoRNAs do interact with non-rRNA transcripts, and these interactions can be captured and validated using an optimized snoCLASH coupled with computational and biochemical approaches. This work expands the functional capacity of snoRNAs and lays the foundation for uncovering new layers of RNA-based regulation with implications for metabolism, stress signaling, and human disease.

## ACKNOWLEDGEMENTS

The authors acknowledge the support of the Freiburg Galaxy Team for use of the ChiRA pipeline: Bjorn Gruning, Bioinformatics, University of Freiburg (Germany), funded by the German Federal Ministry of Education and Research BMFTR grant 031 A538A de.NBI-RBC and the Ministry of Science, Research and the Arts Baden-Württemberg (MWK) within the framework of LIBIS/de.NBI Freiburg.

## AUTHOR CONTRIBUTIONS

B.A.E. and C.L.H. conceived and designed the study. B.A.E. performed the majority of experiments, developed methodology, conducted formal and computational analyses, curated data, validated findings, and wrote the original manuscript draft. G.Y. contributed to methodology development, computational analysis, and resources. A.K.C. contributed to experimentation and methodology. Y.Z. contributed to sequencing data production, curation, formal analysis, and methodology. W.R.F. contributed to investigation and validation experiments. C.L.H. supervised the project and contributed to study conceptualization, methodology, resources, funding acquisition, and manuscript review and editing. All authors reviewed and approved the final manuscript.

## SUPPLEMENTARY DATA

Supplementary Data will be available online at the time of publication.

## DATA AVAILABILITY

Raw data will be available on GEO at the time of publication.

