## Supplemental Figures for "snoCLASH Reveals Extensive snoRNA-mRNA Interaction Networks"

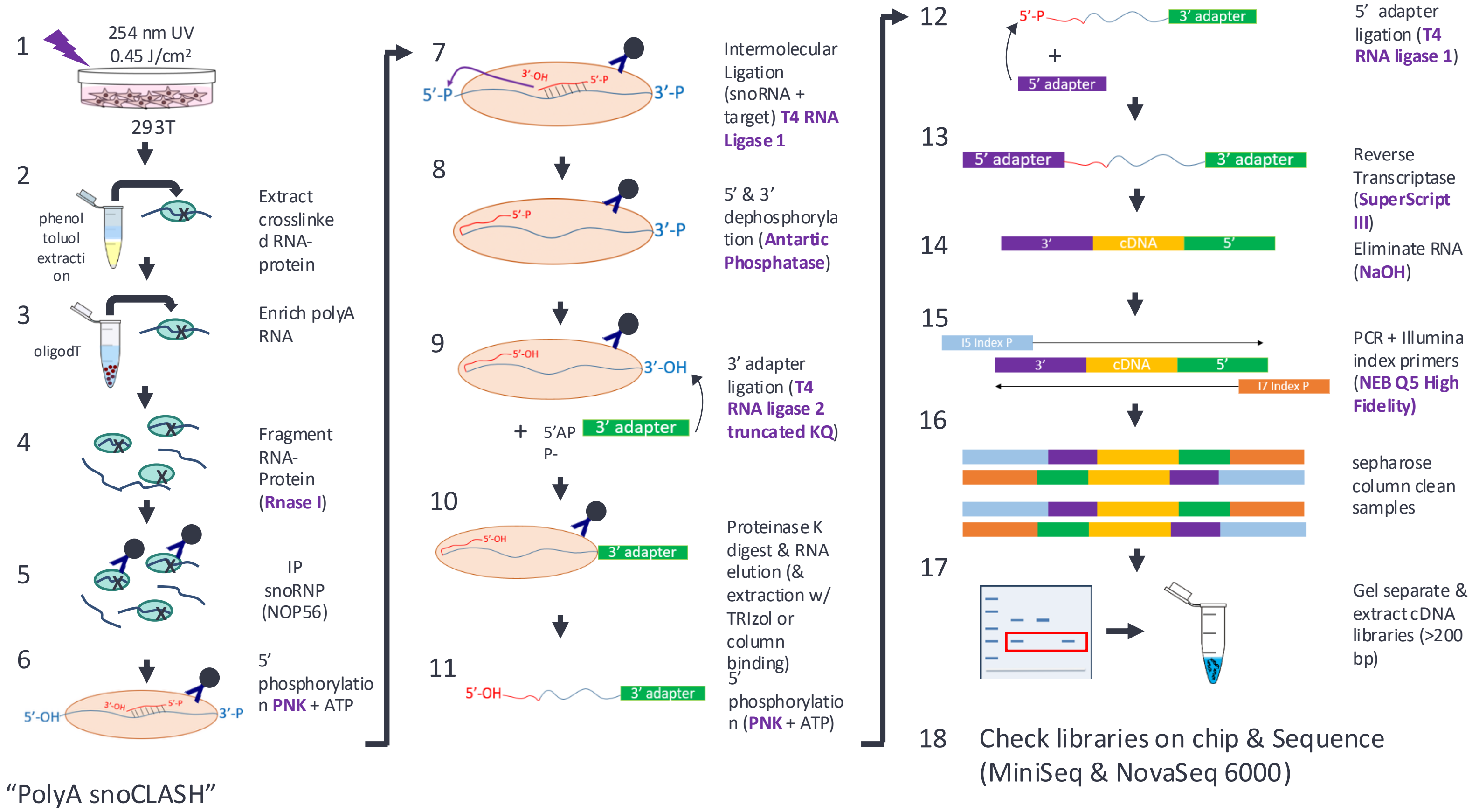


**Supplementary Figure 1. Detailed workflow for PolyA-snoCLASH.** Schematic overview of the PolyA-snoCLASH workflow used to enrich for snoRNA–mRNA interactions. HEK293T cells were UV crosslinked (254 nm, 0.45 J/cm²) to stabilize RNA–protein interactions, followed by phenol–toluol extraction (PTex) to selectively purify crosslinked RNA–protein complexes. Polyadenylated RNA species were enriched using oligo-dT selection prior to immunoprecipitation of snoRNP complexes via NOP56. RNA–protein complexes were fragmented using RNase I and subjected to intermolecular ligation to capture snoRNA–target RNA hybrids. Following ligation, RNA fragments were dephosphorylated and ligated to 3′ and 5′ adapters, and protein was removed by Proteinase K digestion. RNA was extracted and converted to cDNA by reverse transcription, followed by PCR amplification using Illumina-compatible primers. Libraries were size-selected (>200 bp), purified, and sequenced. This workflow enriches for polyadenylated, non-ribosomal RNA targets associated with snoRNAs.

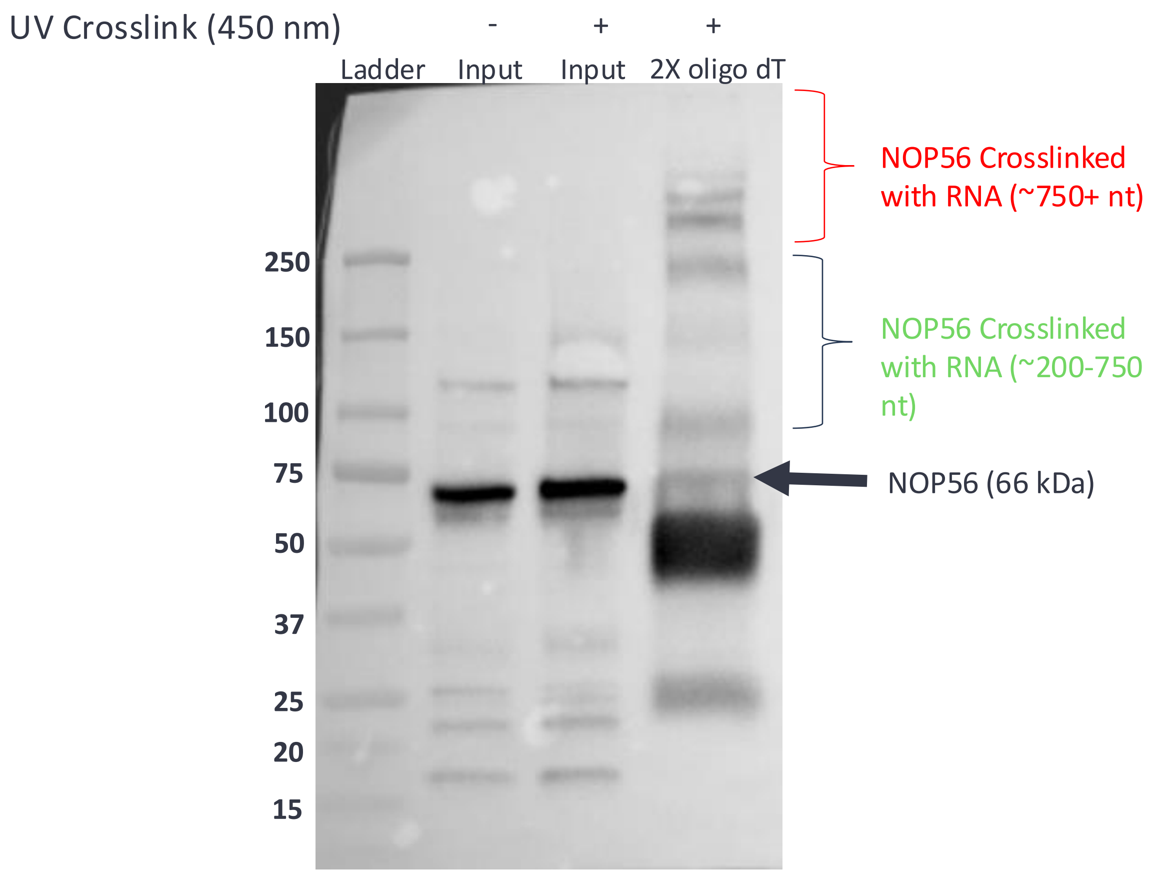


**Supplementary Figure 2. NOP56 IP/Western shows that polyA selection of crosslinked RNA samples captures NOP56-mRNA complexes.** 293T cells were UV crosslinked at 450 nm and RNA-protein (RNP) complexes were extracted based on pH separation (phenol-toluol extraction). Samples were enriched for mRNA by two selections with oligo-dT coated magnetic beads. The poly(A) enriched RNA (mRNA) was then immunoprecipitated for NOP56 using protein A Dynabeads. Blot was probed for NOP56 using the same anti-NOP56 antibody. **Lane 1**: No Crosslink, phenol-toluol extraction; **Lane 2**: 450 nm UV Crosslink, phenol-toluol extraction; **Lane 3**: 450 nm UV crosslink, phenol-toluol extraction, poly(A) enrichment, NOP56 immunoprecipitation.

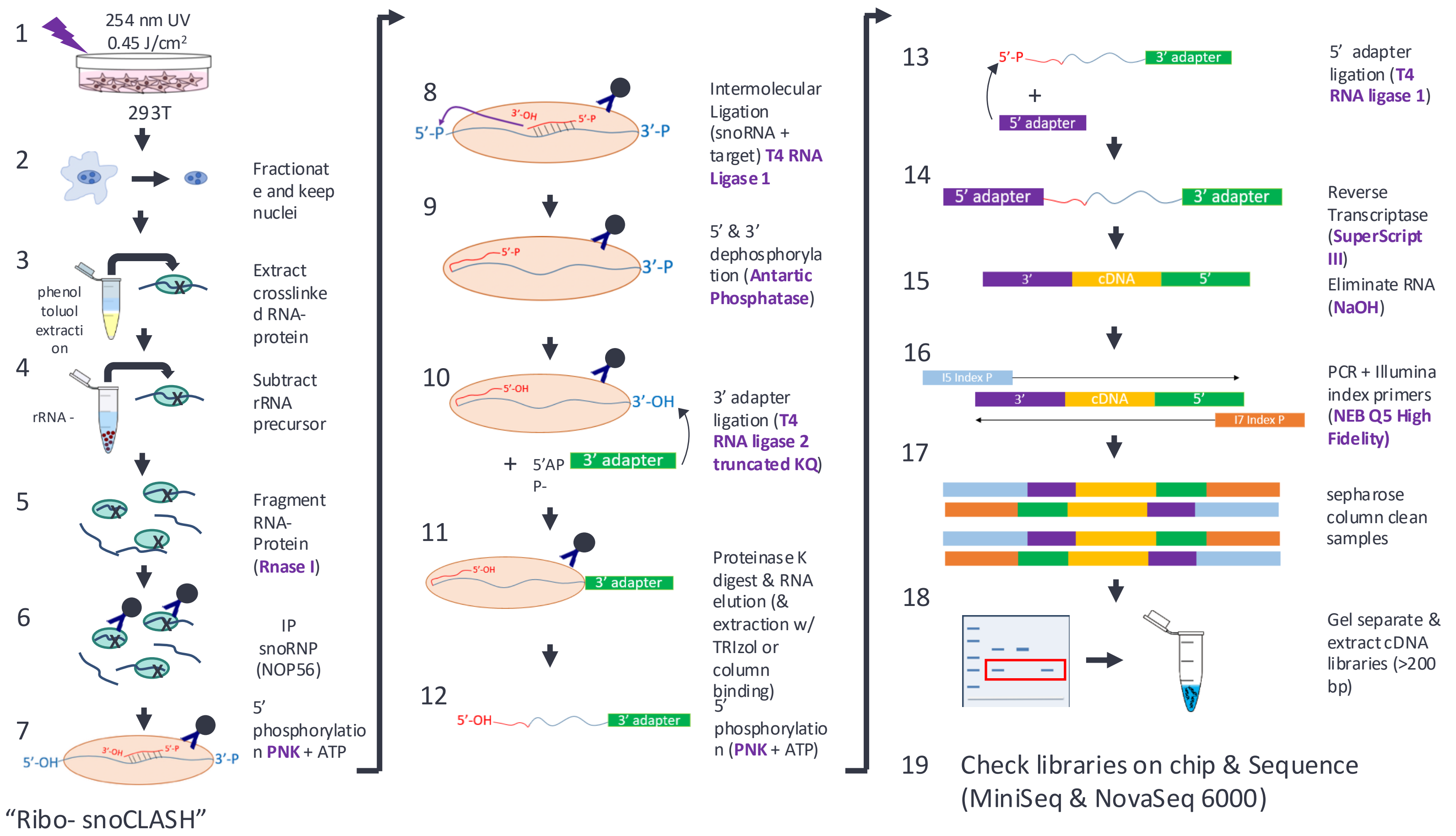


**Supplementary Figure 3. Detailed workflow for Ribo(-)-snoCLASH.** Schematic overview of the Ribo(-)-snoCLASH workflow designed to reduce ribosomal RNA contamination and enrich for nuclear snoRNA–target interactions. HEK293T cells were UV crosslinked (254 nm, 0.45 J/cm²), followed by nuclear fractionation to remove cytoplasmic ribosomes. Crosslinked RNA–protein complexes were purified using phenol–toluol extraction (PTex), and ribosomal RNA and precursor rRNA species were depleted using Ribo(-) reagents. snoRNP complexes were immunoprecipitated via NOP56, and RNA fragments were generated by RNase I digestion. Intermolecular ligation was performed to capture snoRNA–target hybrids, followed by adapter ligation, Proteinase K digestion, and RNA extraction. cDNA libraries were generated by reverse transcription and PCR amplification, size-selected (>200 bp), and sequenced. This workflow enhances detection of non-ribosomal targets, particularly within nuclear RNA populations.

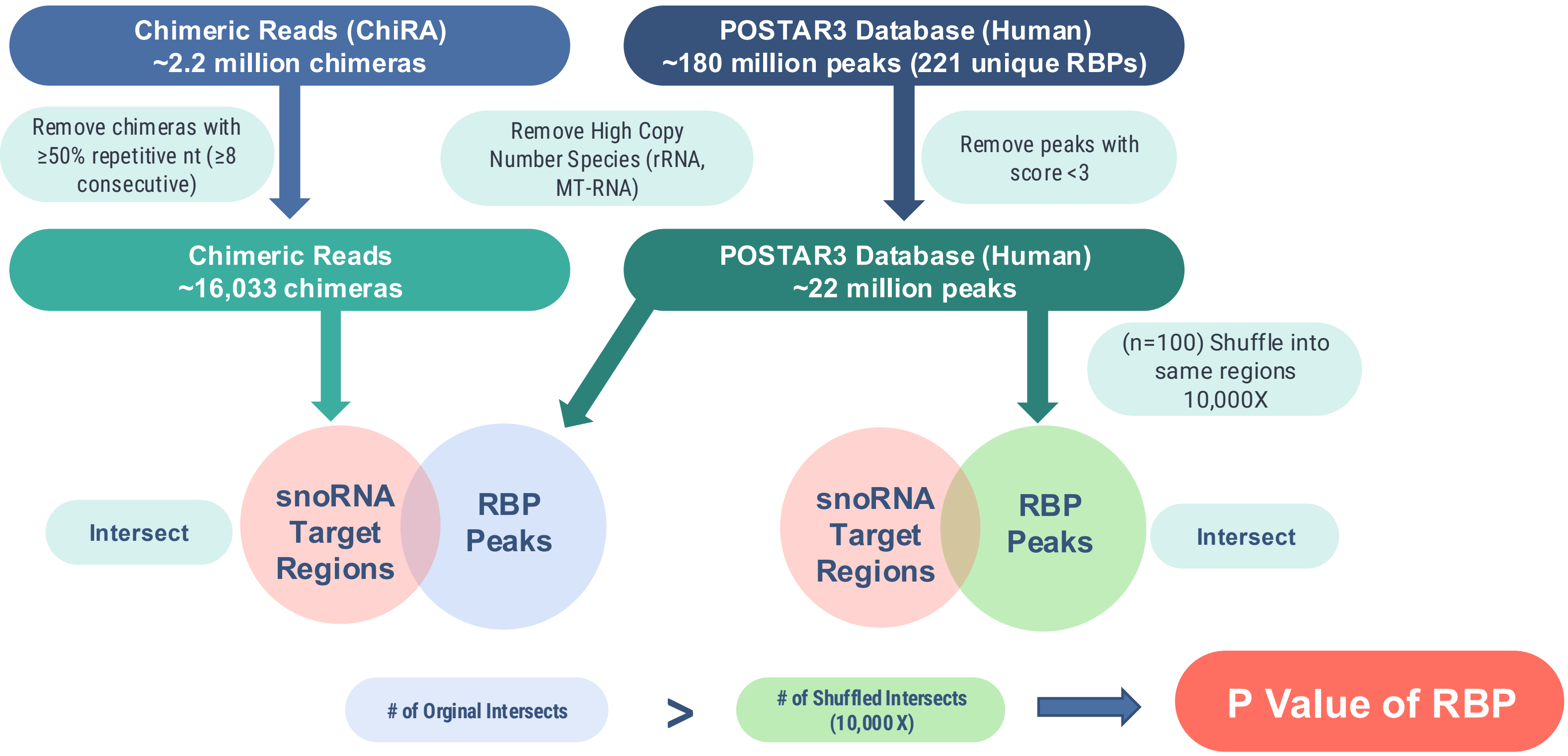


**Supplementary Figure 4. Schematic of snoRNA-target chimera filtering and RBP intersection analysis.** Overview of the computational pipeline used to identify high-confidence snoRNA-target interactions and assess overlap with RNA-binding protein (RBP) binding sites. Raw chimeric reads identified by ChiRA were filtered to remove duplicate sequences, low-complexity reads containing ≥8 consecutive identical nucleotides, and high-copy RNA species including ribosomal and mitochondrial RNAs. The resulting high-confidence snoRNA-target regions were intersected with experimentally derived RBP binding sites from the POSTAR3 database. To assess statistical enrichment, RBP binding sites were randomly shuffled within transcript regions of the same annotation class (e.g., CDS, intron, UTR) for 10,000 iterations and overlap with snoRNA-target regions was recalculated to generate an empirical null distribution. Observed overlaps were compared to this null distribution to identify RBPs significantly enriched at snoRNA-target sites.

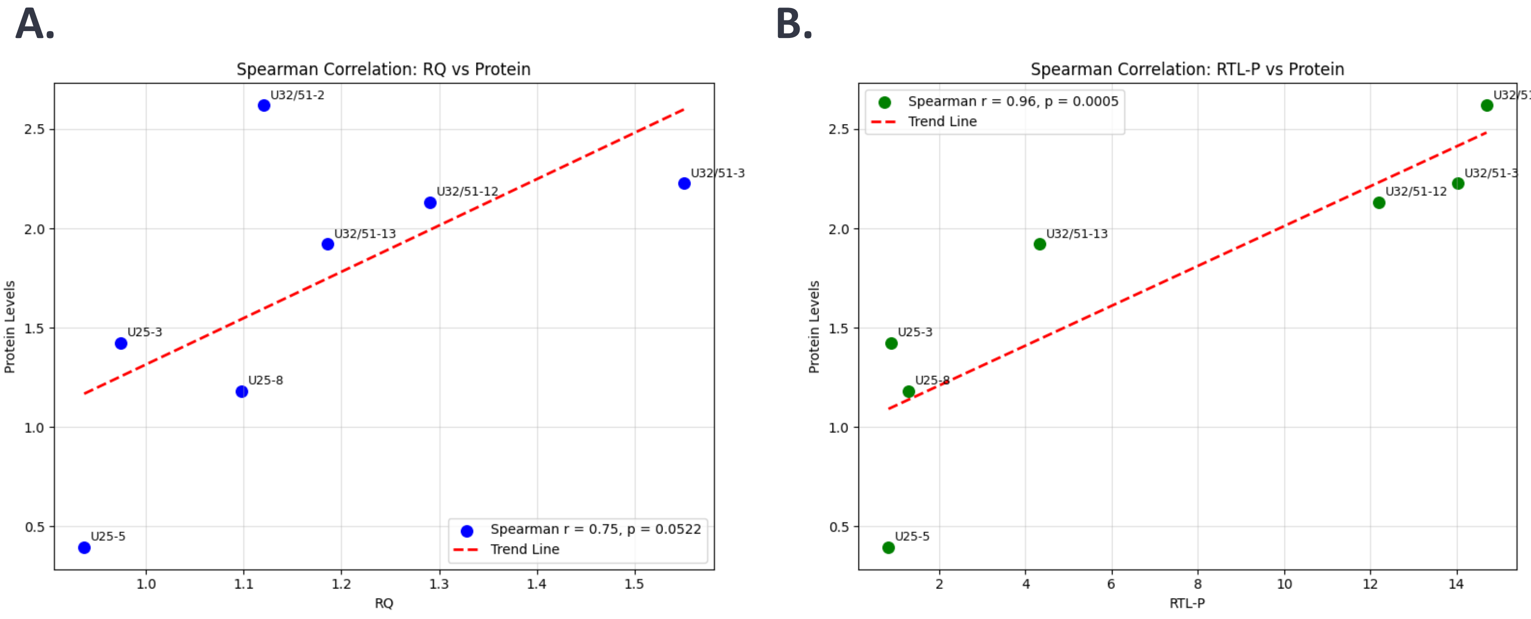


**Supplementary Figure 5. Correlation of transcript abundance and RTL-P signal with OGA protein expression.** Spearman correlation analysis comparing OGA protein expression with **A.** relative transcript abundance (RQ) and **B.** RTL-P–derived measurements of 2′-O-methylation. Correlation coefficients (ρ) were calculated to assess the relationship between mRNA levels and protein output, as well as between methylation-dependent reverse transcription efficiency and protein expression. These analyses evaluate whether changes in OGA protein levels are more strongly associated with transcript abundance or with snoRNA-guided 2′-O-methylation.
